# The RNA-binding protein NOVA-1 regulates circRNA expression, alternative splicing, and aging in *C. elegans*

**DOI:** 10.1101/2025.10.02.679314

**Authors:** Emmanuel Adeyemi, Hussam Z. Alshareef, Jaffar M. Bhat, Pedro Miura, Alexander M. van der Linden

## Abstract

Circular RNA (circRNA) biogenesis is regulated by RNA-binding proteins (RBPs) that alter back-splicing of exons in protein coding genes. However, few *in vivo* roles for RBPs in the regulation of circRNA biogenesis have been characterized. We previously showed that many circRNAs increase with age in *C. elegans*, and that loss of circ-*crh-1*, an abundant age-accumulated circRNA, extends mean lifespan. Given the established role of the mammalian RBP NOVA2 in promoting circRNA biogenesis, we investigated whether *nova-1*, the sole *C. elegans* homolog of NOVA1/2, similarly regulates circRNA expression and function *in vivo*. RNA-sequencing of *nova-1* mutants compared to wild-type identified 686 circRNAs. Of these, 103 were differentially expressed in *nova-1* mutants compared to wild-type, with 76 upregulated and 27 downregulated circRNAs, suggesting NOVA-1 acts as a negative regulator of a subset of circRNAs. *nova-1* mutants also exhibited linear alternative splicing changes, primarily in alternative 3′ splice site usage and exon skipping, and showed minimal overlap with circRNA loci. Notably, circ-*crh-1* represented a shared regulatory target, suggesting NOVA-1 may coordinate splicing regulation with the production of *crh-1* circRNAs. Motif analysis further revealed that over half of the NOVA-1-regulated splicing events contained YCAY motif sites, with *crh-1* harboring a high density of sites, consistent with its alternative 3′ splice site usage and circRNA production. Finally, *nova-1* mutants exhibited an extended mean lifespan and enhanced heat stress recovery. Together, these findings identify NOVA-1 as a key regulator of circRNA expression and alternative splicing in *C. elegans*, with likely downstream consequences for organismal lifespan and stress resilience.

## Introduction

Circular RNAs (circRNAs) are generated through back-splicing, a non-canonical splicing event in which a downstream 5′ splice donor (SD) joins an upstream 3′ splice acceptor (SA), forming a covalently closed loop marked by a unique back-splice junction (BSJ) (Cortes-Lopez and Miura 2016). Back-splicing is regulated by a complex interplay of cis-elements and trans-acting factors that promote looping of intronic regions flanking the circularizing exon(s) (Chen 2020), thereby facilitating circRNA formation. Inverted repeat sequences within introns can base-pair to bring SD and SA sites into proximity (Ashwal-Fluss et al. 2014; Ivanov et al. 2015), while RNA-binding proteins (RBPs) enhance back-splicing by stabilizing looped RNA structures and binding intronic motifs (Conn et al. 2015; Errichelli et al. 2017).

The NOVA family of splicing factors, NOVA1 and NOVA2, have emerged as key regulators of splicing in neuronal tissues (Piton 2025). NOVA2, a neuron-specific splicing factor that binds YCAY motifs (where Y indicates pyrimidines T or C), is essential for promoting circRNA formation during brain development (Knupp et al. 2021). Embryonic Nova2 knockout mice exhibit a global reduction in neuronal circRNAs, both in whole embryonic cortex and in isolated excitatory and inhibitory neurons (Saito et al. 2016; Knupp et al. 2021), underscoring NOVA2’s role in facilitating circRNA formation in cortical neurons. In contrast, NOVA1, a paralog of NOVA2 with functions in alternative splicing and neurons (Buckanovich et al. 1993; Jensen et al. 2000), does not cause a widespread effect on circRNA levels when deleted (Saito et al. 2016; Knupp et al. 2021) indicating that circRNA regulation is a unique function of NOVA2. Notably, NOVA2-regulated circRNAs show minimal overlap with its exon-skipping targets (Knupp et al. 2021), suggesting a mechanistically distinct role for NOVA2 in circRNA biogenesis apart from alternative splicing activity. However, *in vivo* functional analysis of NOVA proteins remains limited due to the early lethality of NOVA-null mice.

The nematode *Caenorhabditis elegans* expresses a single NOVA homolog, *nova-1*, providing a genetically tractable model to investigate the *in vivo* role of NOVA proteins in circRNA regulation and alternative splicing. While mammalian NOVA2 promotes circRNA formation independently of its splicing function (Knupp et al. 2021), it remains unexplored whether *C. elegans nova-1* exerts similar dual functions. Here, we show that *nova-1* regulates circRNA expression and alternative splicing, largely through distinct mechanisms. RNA-seq of *nova-1* mutants revealed 686 differentially expressed circRNAs including the lifespan-associated circ-*crh-1*. Splicing analysis revealed significant changes in alternative 3′ splice site usage and exon skipping, with little overlap between circRNA regulation and splicing targets mediated by NOVA-1.

However, the expression of circ-*crh-1* represents a unique shared regulatory target in which NOVA-1 influences *crh-1* circRNA formation through 3’-alternative splicing. Lastly, *nova-1* mutants have an extended mean lifespan and improved heat stress recovery, highlighting NOVA-1 as a key regulator of aging and stress resilience in *C. elegans*. Together these findings suggest that NOVA-1 is a key regulator of circRNA expression and alternative splicing in *C. elegans*, with likely downstream consequences for organismal lifespan and stress resilience.

## Methods

### *C. elegans* maintenance

Worms were cultivated on the surface of NGM agar seeded with the *Escherichia coli* strain OP50 as the primary food source and grown in 20°C incubators using standard protocols unless indicated otherwise. We used the wild-type strain N2, variety Bristol (Brenner 1974) and VDL1105 *nova-1(tm6146)* mutant strain, which was outcrossed at least six times. The *nova-1* genotype was confirmed by PCR by identifying the 582 bp deletion using specific primers (see **Table S1**).

### Total RNA collection and extraction

Worms were age-synchronized by hypochlorite treatment and collected eggs were hatched overnight at 20°C in 1x M9 buffer. L1 larvae were then plated onto NGM plates seeded with 10x concentrated *E. coli* OP50 bacteria and allowed to develop to the L4 larval stage at 20°C. L4 larvae were then collected, washed, and re-plated onto 25 µM 5-fluorodeoxyuridine (FUdR) (Milipore Sigma, Cat #50-91-9) containing NGM plates unless indicated otherwise seeded with 10x *E. coli* OP50 bacteria in order to prevent progeny formation. Three-day old adult worms were collected by filtering through a 35 µM nylon mesh to remove bacteria (Sefar, Cat #7050-1220-000-10). Worm pellets were then transferred into green RINO tubes (Next Advance) and TRizol LS reagent (ThermoFisher Scientific, Cat #10296028) was added in a 1:3 ratio. Worms were immediately lysed by bead beating them for 5 min using a Bullet Blender Pro Storm (Next Advance). Total RNA was extracted using the Purelink RNA mini-kit, followed by a DNAse I treatment following the manufacturer’s protocol (Ambion, Cat #12183020). RNA was quantified by a Nanodrop. Bioanalyzer or tapestation (Agilent) were used for qualification as needed, and samples were stored at −80°C.

### RNA-Seq and circRNA prediction, mapping and quantification

1 μg total RNA was extracted from five independent biological replicates of wild-type (N2) and *nova-1(tm6146)* (VDL1105) animals, prepared and sequenced on an Illumina HiSeq 6000 system to obtain paired-end 150nt reads at the University of Connecticut genomics center. Raw FASTQ files were aligned to the WBcel235/ce11 reference genome using HISAT2 v2.2.1 (parameters: -no-mixed -no-discordant). Unmapped reads were realigned to the reference genome using BWA v0.7.18-r1243 mem followed by circRNA loci prediction using CIRI2 v2.0.6 (Gao et al. 2018). Predicted circRNAs with junction reads <12 were filtered out. From the predicted circRNA loci, a 200-nt circRNA junction scaffold was built and Bowtie2 v2.5.4 (Langmead and Salzberg 2012) was used for alignment and mapping to raw FASTQ datasets. Duplicate reads were removed after mapping with Picard (http://broadinstitute.github.io/picard) (parameters: MarkDuplicates ASSUME_SORTED = true REMOVE_DUPLICATES = true). Circular RNA counts in each sample were quantified using featureCounts v2.0.1 (Liao et al. 2014). edgeR v4.0.16 was used to calculate Fold-change count vales between *nova-1* and wild-type samples (Chen et al. 2025). CircRNAs were considered significantly upregulated in *nova-1* mutants if log_2_ fold change (log_2_FC) < −0.5 with *P*<0.05, and downregulated if log_2_FC > 0.5 with *P*<0.05. This threshold corresponds to at least a ∼1.4-fold change in expression. Raw FASTQ files were deposited at the NCBI Sequence Read Archive (BioProject: TBD). Individual accession numbers are listed in **Table S2**.

### Multivariate Analysis of Transcript Splicing (rMATS)

For alternative splicing analysis, we applied rMATS to calculate significant skipped exon (SE), alternative 3’ and 5’ splice site usage (A3’SS & A5’SS), mutually exclusive exons (MXE), and retained intron (RI) splicing events in *nova-1* mutants versus wild-type controls. rMATS utilizes a linear mixed model to compute the ‘Percent Spliced In’ (PSI) value by evaluating the splicing levels of alternative/mutually exclusive exons, alternative 3’ or 5’ splice sites, or intron retention in the samples (25480548, 38396040). Briefly, FASTQ files were mapped using STAR 2.7.10a using default parameters and subsequently aligned to the WBcel235/ce11 genome. rMATS-v.4.3.0 was used to discover significant linear alternative splicing events with the default parameters. For post processing, junction count and exon count (*JCEC.txt) output files of all five splicing events were obtained and uploaded into R. To further process the output, the R package maser (Mapping Alternative Splicing Events to pRoteins) was used (https://www.bioconductor.org/packages/devel/bioc/vignettes/maser/inst/doc/Introduction.html). Significant splicing events were selected by applying filtering threshold of FDR ≤ 0.05 and |ΔPSI| ≥ 0.2 to the ‘topEvents’ maser function, thereby selecting for splicing changes that were at least 20% different between *nova-1* mutants and wild-type. The ‘plotTranscripts’ function was used to visualize splicing events in R. Custom scripts were used to count the number of *nova-1* YCAY ([C|T]CA[C|T]) binding sites in exons of alternatively spliced transcripts.

### RT-qPCR analysis

To quantify and confirm individual circular or linear transcripts, 0.5 μg total RNA was reverse transcribed using Superscript III to prepare cDNA using random hexamers (Invitrogen, Cat #18080051). Next, cDNA samples were diluted and used with PowerUp SYBR Green Master Mix (Applied Biosystems, Cat #A25471) for RT-qPCR analysis analyzed on a CFX96 Real-Time System (Bio-Rad). For RT-qPCRs of circRNAs, we used outward-facing primers. For host gene linear RNA counterparts, one primer was located in the circularizing exon, and the other was located in the upstream or downstream non-circularizing exon. For linear mRNAs, we used forward-facing primers. Fold-change values were calculated using wild-type (N2) ΔCt as control values for the 2^-ΔΔCt^ method. Data is normalized to housekeeping genes (*cdc-42*) mRNA. Primer sequences are listed in **Table S1**.

### Lifespan assay

Strains were maintained at 20°C for at least two generations before the lifespan assay. Adult worms age-synchronized by hypochlorite treatment were allowed to lay eggs on NGM plates seeded with 10x concentrated *E. coli* OP50 bacteria over ∼3 hours, and then removed. The resulting progeny synchronized by the timed egg-laying were allowed to develop into L4 larvae at 20°C. At the L4 stage, 150 worms per genotype were transferred to new 6 cm NGM plates with freshly seeded 10x concentrated *E. coli* OP50 bacteria. Wild-type controls (N2) were assayed in parallel to mutants in the absence of FUdR and at 20°C. Adult worms were transferred every 2 days during active reproduction, and each plate contained 10-15 worms. Worms that experienced ventral rupture, bagging, or walling were censored from the life-span analysis. A worm was considered dead when it did not respond to touch of the platinum wire pick, and was subsequently removed from the plate.

### Heat shock viability assay

Animals age-synchronized by hypochlorite treatment were allowed to grow at 20°C to reach L4 developmental stage. L4 animals were exposed to 35°C heat-shock in a water-bath for 4 hrs. Following the heat shock treatment, worms were recovered at 20°C for 24 hrs on NGM plates seeded with *E. coli* OP50 before scoring viability.

## Results

### NOVA-1 regulates a subset of circRNAs

Using RNA-sequencing, we analyzed circRNA expression in whole *nova-1* mutants compared to whole wild-type worms in 3-day old adults. CircRNA loci were identified from the RNA-seq dataset using CIRI2, a tool that detects back-splice junctions (BSJ) from RNA-seq data (Gao et al. 2018). In total, we detected 686 circRNAs (**Table S1)**. Of these, 103 circRNAs were differentially expressed in *nova-1* mutants compared to wild-type, with 76 upregulated and 27 downregulated circRNAs (**Fig. 1A, Table S1**) (|Fold-change| >1.4, adj. *P* < 0.05). RT-qPCR validation of select circRNAs using outward facing primers confirmed the directionality of expression changes in six selected circRNAs, including three upregulated and three downregulated circRNAs (**Fig. 1B**). Importantly, the corresponding linear transcripts of the host genes did not show significant expression changes (**Fig. 1B**). Notably, 73 of the 103 differentially expressed circRNAs overlapped with a previously characterized set of age-associated circRNAs (Cortés-López et al. 2018). This included *cel_circ_new_0000088*, derived from the *Y52B11A.8* gene, which was significantly downregulated in *nova-1* mutants (*P*= 8.42e-05), suggesting that NOVA-1 regulates the expression of circRNAs enriched during aging.

**Fig. 1:**
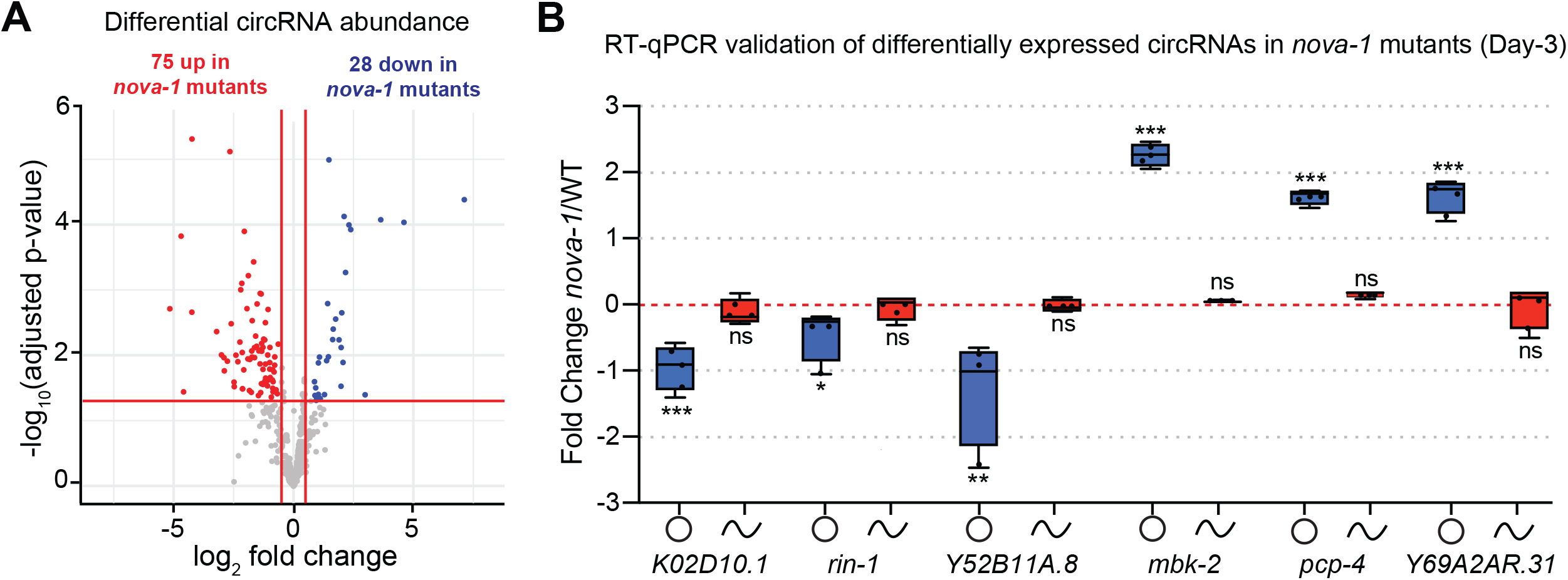
NOVA-1 regulates a subset of circRNAs. **(A)** Volcano plot of differentially expressed circRNAs in whole animals of *nova-1* mutants compared to wild-type control as determined by RNA-seq. Significantly downregulated and upregulated genes (Log_2_ fold-change >2, adj. *P* < 0.05), blue color represents down-regulated circRNAs, red color represents upregulated circRNAs. **(B)** RT-qPCR analysis of circular and linear differentially regulated transcripts in day-3 adults of *nova-1* mutants compared to wild-type. Shown are three upregulated and three downregulated circRNAs. n=3 independent biological replicates. Data was normalized to *cdc-24* mRNA, and is represented as the mean ± SEM; ns, not significant; \*\*\*\**P* < 0.001,\*\*\**P* < 0.01, \*\**P* < 0.05, and \**P* < 0.1.

### Distinct circ-*crh-1* isoform regulation by NOVA-1

We previously identified two circRNAs derived from the *crh-1* gene (i.e., *cel_circ_0000438* and *cel_circ_0000439*) (Knupp et al. 2022). These circRNAs are two of the most abundant, age-accumulated circRNA isoforms derived from the *crh-1* host gene. The circRNAs differ by only in six nucleotides, as a result of an alternative splice acceptor site (A3’SS) in exon 4 of the *crh-1* locus, and are collectively referred to as circ-*crh-1* (**Fig. 2A**). To determine whether NOVA-1 influences circ-*crh-1* formation, we quantified the expression of the two circ-*crh-1* isoforms using RT-qPCR in 3-day old adults. We observed a significant reduction in *cel_circ_0000438* in *nova-1* mutants, while levels of *cel_circ_0000439* and linear *crh-1* remained unchanged (**Fig. 2B**). These findings suggest that loss of NOVA-1 modulates the expression of one circ-*crh-1* isoform (i.e. *cel_circ_0000438*) likely by selectively impairing back-splicing at one acceptor site without affecting the other circ-*crh-1* isoform or linear *crh-1*.

**Fig. 2:**
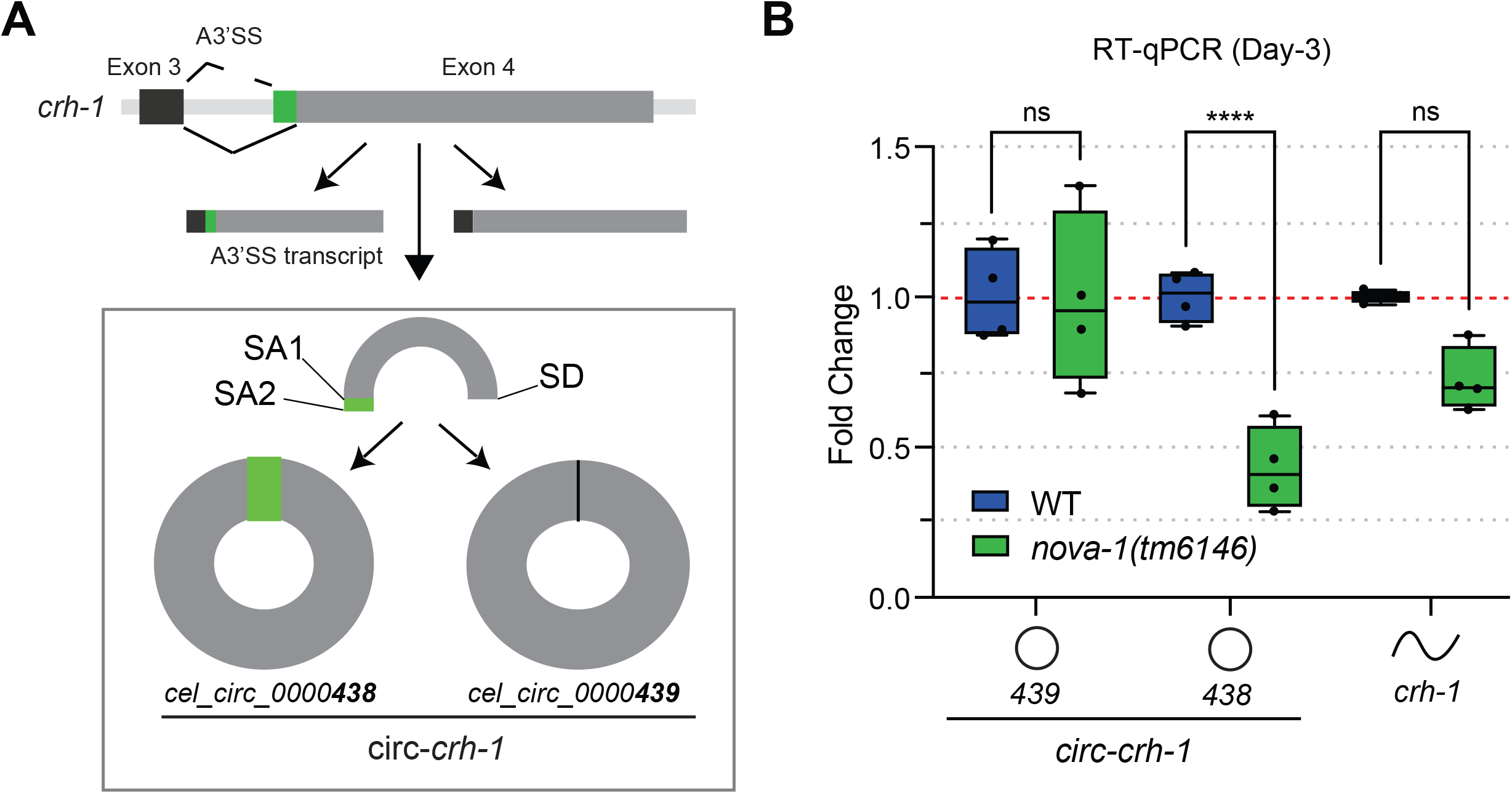
NOVA-1 differentially regulates two circ-*crh-1* isoforms. **(A)** Schematic showing the alternative site usage in exon 4 of *crh-1*. Two circRNAs (*cel_circ_0000438* and *cel_circ_0000439*) are generated by back-splicing of exon 4, using two alternative splice acceptors (SA) and one shared splice donor (SD). **(B)** RT-qPCR analysis of circular and linear circ-*crh-1* transcripts in 3-day old adults at 20°C of *nova-1* mutants compared to wild-type. *nova-1* mutants reduce *cel_circ_0000438* expression but not *cel_circ_0000439*, whereas the linear *crh-1* transcript is not significantly changed. n=4 independent biological samples. Data was normalized to *cdc-24* mRNA and is represented as the mean ± SEM; ns, not significant; \*\*\*\**P* < 0.001.

### NOVA-1 regulates alternative splicing events

We performed analysis of linear alternative splicing using rMATS-turbo v4.1.2 (Shen et al. 2014; Wang et al. 2024) on all wild-type and *nova-1* mutant samples, identifying over 16,000 linear splicing events. After applying stringent filtering (FDR ≤ 0.05, |ΔPSI| ≥ 0.2), 307 total events were classified as differentially expressed in *nova-1* mutants compared to wild-type controls (**Fig. 3A, S1A, Table S4**). Of these, 195 (64%) involved alternative 3′ splice site (A3’SS) usage, and 80 (26%) were skipped exons (SE) (**Fig. 3B, S1A**). The remaining 10% comprised alternative 5′ splice sites (5A’SS), retained introns (RI), and mutually exclusive exons (MXE) (**Fig. 3B, S1A**), indicating widespread splicing disruption in the absence of NOVA-1.

**Fig. 3:**
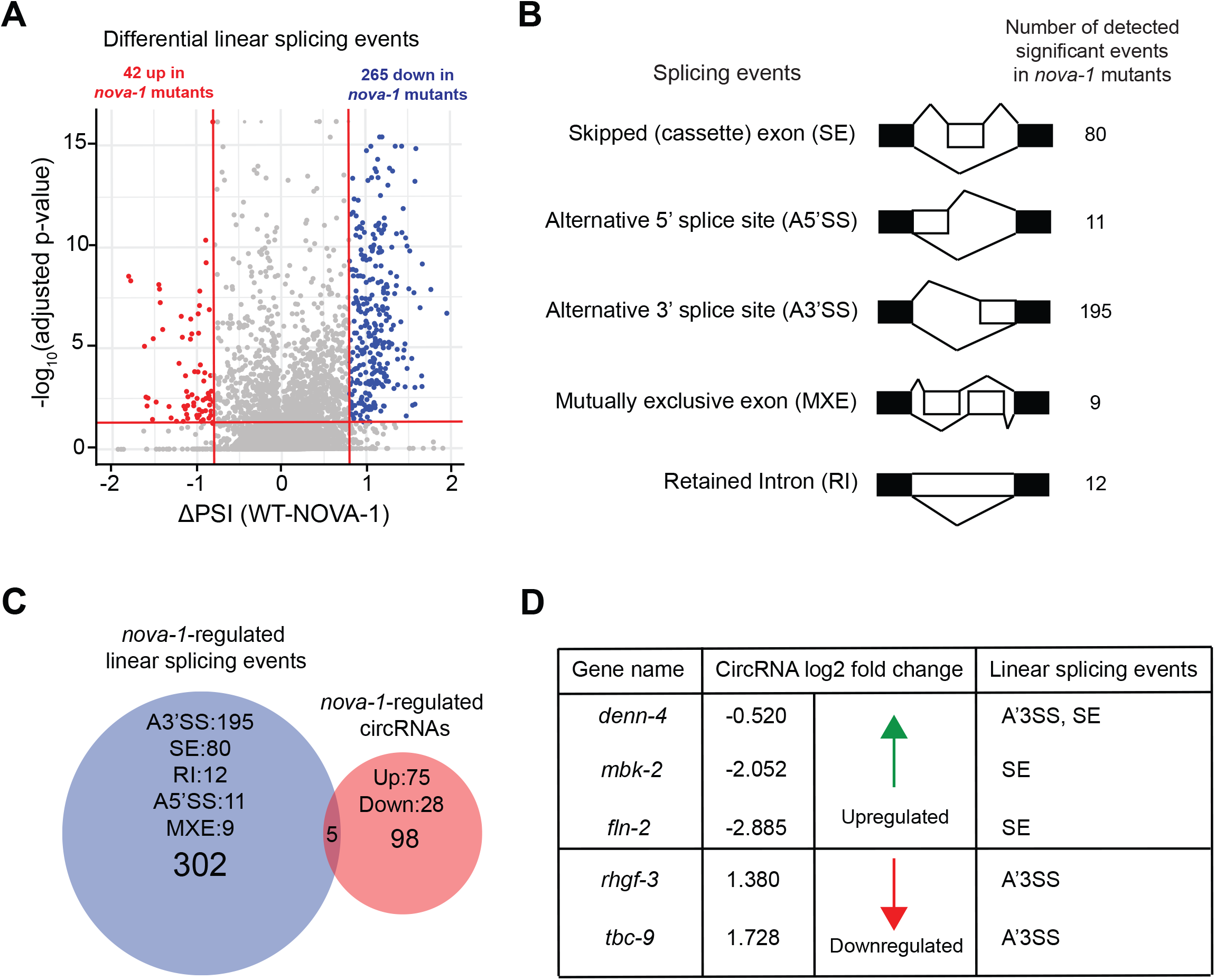
NOVA-1 regulates distinct gene sets during pre-mRNA alternative splicing and back-splicing programs. **(A)** Volcano plot showing total splicing changes in *nova-1* mutants compared to wild-type control. Points represent transcripts from any of the five linear splicing events (i.e., A3’SS, A5’SS, SE, MXE and RI). Upregulated and downregulated splicing events correspond to points with Percent Splice In changes (ΔPSI= WT_PSI – NOVA-1_PSI) ≤ −0.2 (red) and ΔPSI ≥ 0.2 (blue), respectively (FDR ≤ 0.05; adj. *P* < 0.05). **(B)** Schematic representation of the five splicing events and the number of detected significant events in *nova-1* mutants vs. wild-type (FDR ≤ 0.05, |ΔPSI| ≥ 0.2). Alternative 3’ splice site usage (A3’SS) and Skipped exons (SE) are the most abundant of the AS events, both accounting for approximately 90% of *nova-1*-regulated linear splicing events. Alternative 5’ splice site usage (A5’SS), retained introns (RI) and exclusive exons (MXE) account make up the remaining 10% of splicing events regulated by *nova-1*. White exons (boxes) indicate alternatively “spliced in” exons during linear pre-mRNA alternative splicing. **(C-D)** Venn diagram showing the relationship between *nova-1*-regulated linear splicing and back-splicing events. CircRNAs derived from three of the five host genes in this category are more abundant in *nova-1* mutants compared to wild-type controls and all have lower skipped exon PSI (see **D**), with the exception of *denn-4* which, in addition, has a lower A3’SS usage compared to control alongside two *nova-1*-regulated low abundance circRNAs (see **D**).

Next, we examined YCAY motif sites in alternatively spliced transcripts regulated by NOVA-1. Exonic YCAY sites were categorized into three different groups: low (<20 sites), intermediate (20-59 sites), and enriched (>60 sites). This analysis revealed that 168 (55%) of the 307 significant splicing events contained 20 or more YCAY motif sites (**Table S4, S1B**). Focusing on the *crh-1* gene, we identified a total of 43 YCAY sites (**Fig. S1C, Table S4**) with 8 located in exon 4 (**Fig. S1C**), which is subject to A3’SS usage. Notably, this represents at least 3 more YCAY sites than was identified in any other *crh-1* exon, suggesting preferential NOVA-1 binding near exon 4. In addition, 7 and 8 YCAY sites were identified in the upstream and downstream intronic regions flanking exon 4, respectively (**Fig. S1C**). Together, these findings suggest that NOVA-1-mediated alternative splicing may intersect with its role in back-splicing.

### Minimal overlap between NOVA-1-regulated circRNAs and alternative splicing targets

Since *nova-1* mutations affect both back-splicing (103 circRNAs) and alternative splicing events (307 linear transcripts), we next examined potential overlap between these two regulatory processes. Of the 103 differentially expressed circRNAs, only 5 showed overlap with *nova-1*-regulated linear splicing events (**Fig. 3C**). Specifically, 3 of the 75 upregulated and 2 of the 28 downregulated circRNAs overlapped with *nova-1*-regulated linear splicing events (**Fig. 3D**). All overlapping splicing events fell within alternative 3′ splice site (A3’SS) or skipped exon (SE) categories with *denn-4* undergoing both A3’SS usage and SE (**Fig. 3D**). Interestingly, the *crh-1* gene displayed a reduced Percent Spliced In (PSI) at exon 4 (A3’SS) in *nova-1* mutants (**Fig. S1D**). This together with our RT-qPCR analysis (**Fig. 2B**) supports a model in which NOVA-1 differentially regulates the expression of circ-*crh-1* isoforms by modulating alternative splicing. Overall, although NOVA-1 impacts both alternative splicing and circRNA expression, their regulatory overlap is confined to a small subset of targets.

### Loss of NOVA-1 extends mean lifespan and enhanced stress resistance

Given that we previously showed that loss of circ-*crh-1* circRNAs extends mean lifespan (Knupp et al. 2022), we next tested whether *nova-1* mutants exhibit similar lifespan phenotypes. Lifespan assays revealed that *nova-1* mutant animals have a significant 14.75% increase in mean lifespan compared to wild-type controls (*P* < 0.0001, **Fig. 4A**). Because enhanced longevity is often associated with improved stress resistance (Zhou et al. 2011), we also examined thermotolerance as a potential contributing factor. Following a 4-hour heat shock at 35°C, *nova-1* mutant adults showed markedly improved recovery relative to wild-type animals (**Fig. 4B**), suggesting that increased stress resilience may underlie the extended lifespan observed in these *nova-1* mutants. Together these findings suggest that the altered expression of circRNAs together with changes in 3′ alternative splicing and exon skipping events may contribute to the increased mean lifespan and stress resistance phenotype of *nova-1* mutants.

**Fig. 4:**
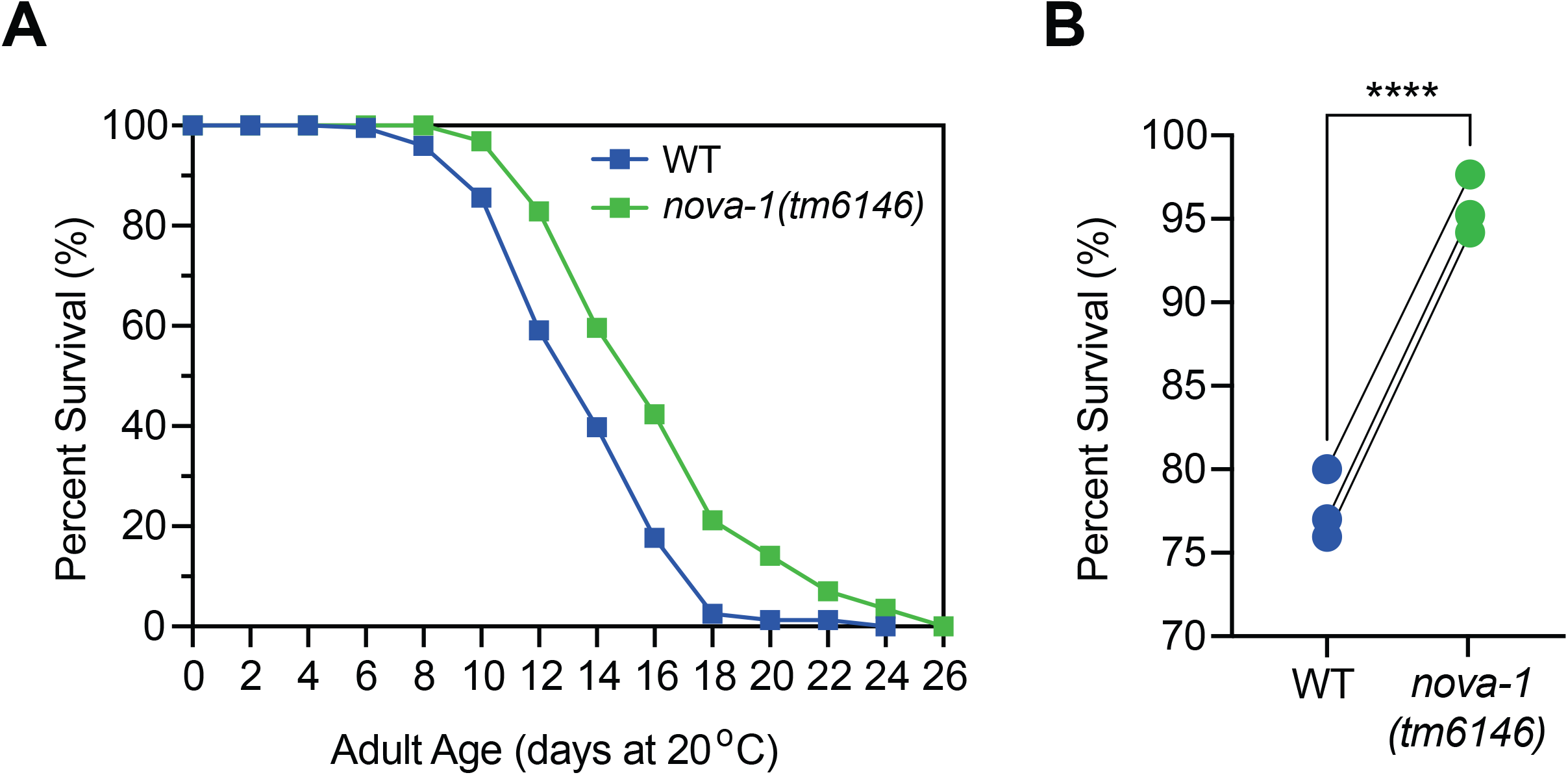
*nova-1* mutants extend mean lifespan and increase heat resistance. **(A)** Lifespan curves for *nova-1* mutant animals compared to wild-type. *nova-1* mutants extend mean lifespan compared to wild-type (*P* < 0.0001, Mantel-Cox log-rank test). n=2 independent lifespan assays were performed with n=150 animals at 20°C for each assay and genotype in the absence of FUdR. **(B)** Loss of *nova-1* during heat shock results in increased viability. Heat shock treatment was performed on synchronized L4 worms for 4 hours at 35°C and the percentage of survival was determined after 24 hrs recovery at 20°C for *nova-1* mutants compared to wild-type controls. *nova-1* mutants exhibit increased resistance to heat exposure compared to wild-type (\*\*\*\**P* < 0.001, Unpaired *t*-test). 3 biological replicates with n=90 animals for each survival assay and genotype in the absence of FUdR.

## Discussion

Our study shows that the RNA-binding protein NOVA-1 regulates the expression of a subset of circRNAs in *C. elegans*, including the age-associated *cel_circ_0000438* (circ-*crh-1*). Using RNA-seq and RT-qPCR analysis, we identified circRNAs whose expression is altered in *nova-1* mutants. Interestingly, the majority of differentially expressed circRNAs were upregulated in *nova-1* mutants, suggesting that unlike NOVA2 in mammals where it promotes circRNA production by binding intronic YCAY motifs to enhance back-splicing in neural tissues (Knupp et al. 2021), NOVA-1 in *C. elegans* may act as a negative regulator of circRNA expression. However, because our RNA-seq analysis was performed on whole worms, it remains possible that NOVA-1 in *C. elegans* promotes circRNA formation in select tissues such as neurons.

Beyond circRNA regulation, we observed widespread splicing disruptions in *nova-1* with the majority of significant changes involving alternative 3′ splice site (A3’SS) selection and exon skipping (SE). This enrichment of A3’SS is consistent with findings from mammalian systems where it was observed that NOVA1/2 proteins bind YCAY sites to regulate diverse alternative splicing outcomes including cassette exons, intron retention and importantly A3’SS usage (Ule et al. 2005; Licatalosi et al. 2008; Zhang et al. 2010). More recently, over 2,000 NOVA1/2-regulated splicing events were identified in the developing mouse cortex, many of which involved A3’SS usage (Saito et al. 2016). Thus, the enrichment of A3’SS events in *C. elegans nova-1* mutants is highly similar with NOVA’s established splicing functions in vertebrates. Moreover, similar to mammalian NOVA2, where circRNA abundance is largely uncoupled from linear alternative splicing (Knupp et al. 2021), we observed that only a small subset of differentially expressed circRNAs in *nova-1* mutants overlapped with genes exhibiting altered linear splicing. The *crh-1* locus illustrates this partial convergence where reduced expression of *cel_circ_0000438* in *nova-1* mutants corresponds with a shift in splice acceptor site usage, whereas expression of the closely related *cel_circ_0000439* and linear *crh-1* remain unaffected. These findings suggests that NOVA-1 in *C. elegans* functions through both shared mechanisms by modulating splice acceptor site choice that influences circRNA back-splicing and distinct pathways that regulate circRNA levels independent of linear splicing.

The position of YCAY motif sites in pre-mRNA determines how NOVA influences splicing regulation (Ule et al. 2006; Witten and Ule 2011). Additionally, NOVA selectively inhibits or promotes alternative splicing regulation depending on whether the sites are exonic or intronic, respectively (Ule et al. 2006). We identified YCAY sites in exon sequences and found enrichment of these sites in over half (55%) of the NOVA-1-regulated transcripts. Interestingly, this exonic enrichment did not result in widespread inhibition of NOVA-1-regulated alternative splicing events in wild-type *C. elegans*, as observed in mammals (Ule et al. 2006). A reasonable explanation may be that *C. elegans* introns are generally shorter and lack strong polypyrimidine tracts compared to mammals, which could alter the way NOVA-1 engages with the spliceosome and reduce the likelihood of reproducing the same splicing regulatory outcomes as observed in mammals. Despite this apparent difference, our analysis revealed that SE still represented the most affected alternative splicing category, with 24 of 80 events (30%) reduced in wild-type compared to *nova-1* mutants. Overall, while the NOVA-mediated bidirectional control of alternative splicing established in mammals provides useful general insights (Ule et al. 2006), our findings suggest that NOVA-regulated alternative splicing in *C. elegans* follows the same underlying principles but plays out differently due to species-specific intron differences and splicing constraints.

Taken together, these findings extend our view of NOVA proteins as key regulators of alternative splicing and circRNA regulation. This study expands our understanding of RBP-mediated circRNA regulation and highlights NOVA-1’s dual role in circRNA expression and splice site choice that may contribute to the regulation of lifespan and stress resistance in *C. elegans*.

## Data Availability

Worm strains are available from the authors. All data necessary for confirming the conclusions are present within the article, figures and tables.

## Acknowledgements

We thank members of the Van Der Linden and Miura labs for useful discussion and feedback on the manuscript. We thank the UConn Genomics Center for performing RNA-sequencing. We thank Thomas Parchman for kindly providing the computational resources for RNA-Seq analysis. We also thank the Cellular and Molecular Imaging Core facility of the COBRE Integrative Neuroscience Center at the University of Nevada for providing equipment and resources. We thank the National BioResource Project (NRBP) for *C. elegans*, funded by the Japanese government for generating and providing the *nova-1(tm6146)* mutant strain.

## Figure Legends

**Fig. S1: NOVA-1 modulates alternative splice site selection at the *crh-1* locus. (A)** Venn diagram of five splicing events with the number of detected significant events in *nova-1* mutants vs. wild-type (FDR ≤ 0.05, |ΔPSI| ≥ 0.2) for alternative 3’ splice site usage (A3’SS), alternative 5’ splice site usage (A5’SS), skipped exons (SE), retained introns (RI), and exclusive exons (MXE). **(B)** Number of exonic YCAY motif sites within alternatively spliced transcripts regulated by NOVA-1 categorized by the number of YCAY sites: low (<20), intermediate (20–59), and enriched (>60). **(C)** Schematic of the *crh-1* gene with a total of 43 YCAY motif sites with 8 shown in exon 4, which is subject to A3’SS usage. In addition, 7 and 8 YCAY sites are shown in upstream and downstream intronic sequences of exon 4, respectively. **(D)** Reduced Percent Spliced In (PSI) at exon 4 of *crh-1* (A3’SS) in *nova-1* mutants.

**Table S1: Oligonucleotide primers used in this study**.

**Table S2: RNA sequencing read statistics**

**Table S3: Detected and differentially expressed circRNAs**

**Table S4: Differential linear splicing events**

## Literature cited

Ashwal-Fluss R, Meyer M, Pamudurti NR, Ivanov A, Bartok O, Hanan M, Evantal N, Memczak S, Rajewsky N, Kadener S. 2014. circRNA biogenesis competes with pre-mRNA splicing. Mol Cell 56: 55–66.

Brenner S. 1974. The genetics of Caenorhabditis elegans. Genetics 77: 71–94.

Buckanovich RJ, Posner JB, Darnell RB. 1993. Nova, the paraneoplastic Ri antigen, is homologous to an RNA-binding protein and is specifically expressed in the developing motor system. Neuron 11: 657–672.

Chen LL. 2020. The expanding regulatory mechanisms and cellular functions of circular RNAs. Nat Rev Mol Cell Biol 21: 475–490.

Chen Y, Chen L, Lun ATL, Baldoni PL, Smyth GK. 2025. edgeR v4: powerful differential analysis of sequencing data with expanded functionality and improved support for small counts and larger datasets. Nucleic Acids Res 53.

Conn SJ, Pillman KA, Toubia J, Conn VM, Salmanidis M, Phillips CA, Roslan S, Schreiber AW, Gregory PA, Goodall GJ. 2015. The RNA binding protein quaking regulates formation of circRNAs. Cell 160: 1125–1134.

Cortés-López M, Gruner MR, Cooper DA, Gruner HN, Voda AI, van der Linden AM, Miura P. 2018. Global accumulation of circRNAs during aging in Caenorhabditis elegans. BMC Genomics 19: 8.

Cortes-Lopez M, Miura P. 2016. Emerging Functions of Circular RNAs. Yale J Biol Med 89: 527–537.

Errichelli L, Dini Modigliani S, Laneve P, Colantoni A, Legnini I, Capauto D, Rosa A, De Santis R, Scarfo R, Peruzzi G et al. 2017. FUS affects circular RNA expression in murine embryonic stem cell-derived motor neurons. Nat Commun 8: 14741.

Gao Y, Zhang J, Zhao F. 2018. Circular RNA identification based on multiple seed matching. Brief Bioinform 19: 803–810.

Ivanov A, Memczak S, Wyler E, Torti F, Porath HT, Orejuela MR, Piechotta M, Levanon EY, Landthaler M, Dieterich C et al. 2015. Analysis of intron sequences reveals hallmarks of circular RNA biogenesis in animals. Cell Rep 10: 170–177.

Jensen KB, Dredge BK, Stefani G, Zhong R, Buckanovich RJ, Okano HJ, Yang YY, Darnell RB. 2000. Nova-1 regulates neuron-specific alternative splicing and is essential for neuronal viability. Neuron 25: 359–371.

Knupp D, Cooper DA, Saito Y, Darnell RB, Miura P. 2021. NOVA2 regulates neural circRNA biogenesis. Nucleic Acids Res 49: 6849–6862.

Knupp D, Jorgensen BG, Alshareef HZ, Bhat JM, Grubbs JJ, Miura P, van der Linden AM. 2022. Loss of circRNAs from the crh-1 gene extends the mean lifespan in Caenorhabditis elegans. Aging Cell 21: e13560.

Langmead B, Salzberg SL. 2012. Fast gapped-read alignment with Bowtie 2. Nat Methods 9: 357–359.

Liao Y, Smyth GK, Shi W. 2014. featureCounts: an efficient general purpose program for assigning sequence reads to genomic features. Bioinformatics 30: 923–930.

Licatalosi DD, Mele A, Fak JJ, Ule J, Kayikci M, Chi SW, Clark TA, Schweitzer AC, Blume JE, Wang X et al. 2008. HITS-CLIP yields genome-wide insights into brain alternative RNA processing. Nature 456: 464–469.

Piton A. 2025. NOVA1/2 genes and alternative splicing in neurodevelopment. Curr Opin Genet Dev 93: 102373.

Saito Y, Miranda-Rottmann S, Ruggiu M, Park CY, Fak JJ, Zhong R, Duncan JS, Fabella BA, Junge HJ, Chen Z et al. 2016. NOVA2-mediated RNA regulation is required for axonal pathfinding during development. Elife 5.

Shen S, Park JW, Lu ZX, Lin L, Henry MD, Wu YN, Zhou Q, Xing Y. 2014. rMATS: robust and flexible detection of differential alternative splicing from replicate RNA-Seq data. Proc Natl Acad Sci U S A 111: E5593–5601.

Ule J, Stefani G, Mele A, Ruggiu M, Wang X, Taneri B, Gaasterland T, Blencowe BJ, Darnell RB. 2006. An RNA map predicting Nova-dependent splicing regulation. Nature 444: 580–586.

Ule J, Ule A, Spencer J, Williams A, Hu JS, Cline M, Wang H, Clark T, Fraser C, Ruggiu M et al. 2005. Nova regulates brain-specific splicing to shape the synapse. Nat Genet 37: 844–852.

Wang Y, Xie Z, Kutschera E, Adams JI, Kadash-Edmondson KE, Xing Y. 2024. rMATS-turbo: an efficient and flexible computational tool for alternative splicing analysis of large-scale RNA-seq data. Nat Protoc 19: 1083–1104.

Witten JT, Ule J. 2011. Understanding splicing regulation through RNA splicing maps. Trends Genet 27: 89–97.

Zhang C, Frias MA, Mele A, Ruggiu M, Eom T, Marney CB, Wang H, Licatalosi DD, Fak JJ, Darnell RB. 2010. Integrative modeling defines the Nova splicing-regulatory network and its combinatorial controls. Science 329: 439–443.

Zhou KI, Pincus Z, Slack FJ. 2011. Longevity and stress in Caenorhabditis elegans. Aging (Albany NY) 3: 733–753.

