## Supplementary figures and images for "The RNA-binding protein NOVA-1 regulates circRNA expression, alternative splicing, and aging in *C. elegans*"

### Supplemental Figure 1

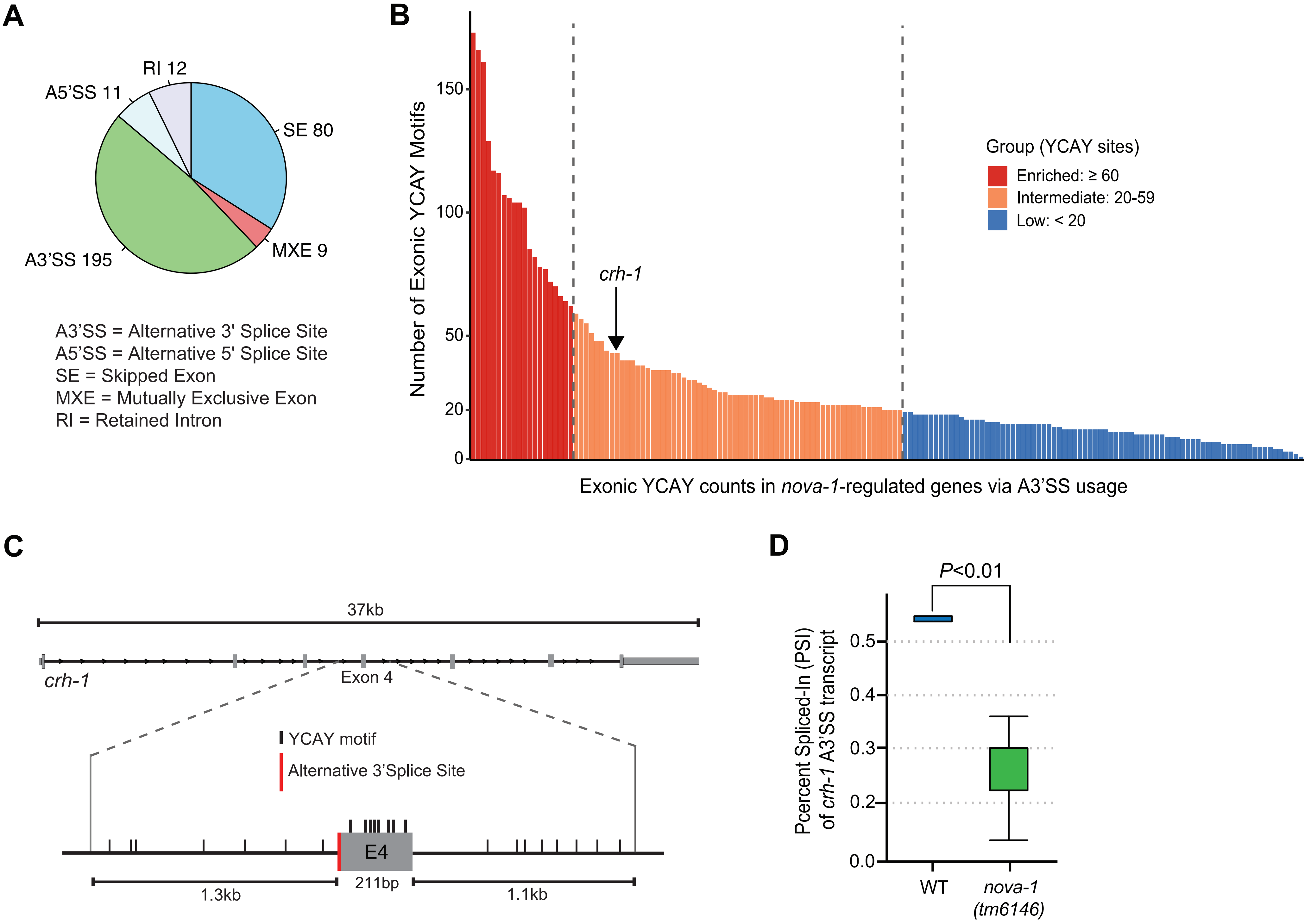
